# Usefulness of the pancreas as a prime target for histopathological diagnosis of *Tilapia parvovirus* (TiPV) infection in Nile tilapia, *Oreochromis niloticus*

**DOI:** 10.1101/2022.03.21.485077

**Authors:** Ha Thanh Dong, Pattiya Sangpo, Le Thanh Dien, Thao Thu Mai, Nguyen Vu Linh, Jorge del-Pozo, Krishna R Salin, Saengchan Senapin

**Author notes:** Corresponding authors: H. T. Dong, S. Senapin.

## Abstract

*Tilapia parvovirus* (TiPV) is an emerging virus reportedly associated with disease and mortality in farmed tilapia. Although previous descriptions of histopathological changes are available, the lesions reported in these are not pathognomonic. Here, we report Cowdry type A inclusion bodies (CAIB) in the pancreas as a diagnostic histopathological feature found in adult Nile tilapia naturally infected with TiPV. This type of inclusion body has been well-known as a histopathological landmark for the diagnosis of other parvoviral infections in shrimp and terrestrial species. Interestingly, this lesion could be exclusively observed in pancreatic acinar cells, both in the hepatopancreas and pancreatic tissue along the intestine. *In situ* hybridization (ISH) using a TiPV-specific probe revealed the intranuclear presence of TiPV DNA in multiple tissues, including the liver, pancreas, kidney, spleen, gills, and the membrane of oocytes in the ovary. These findings suggest that although TiPV can replicate in several tissue types, CAIB manifest exclusively in pancreatic tissues. In addition to TiPV, most diseased fish were co-infected with *Streptococcus agalactiae*, and presented with multifocal granulomas secondary to this bacterial infection. Partial genome amplification of TiPV was successful and revealed high nucleotide identity (> 99%) to previously reported isolates. In summary, this study highlights the usefulness of pancreatic tissue as a prime target for histopathological diagnosis of TiPV in diseased Nile tilapia. This pattern may be critical when determining the presence of TiPV infection in new geographic areas, where ancillary testing may not be available. TiPV pathogenesis in this landmark organ warrants further investigation.

## Introduction

Tilapia (*Oreochromis* spp.) is an aquatic animal species widely cultured in over 100 countries, where global production is approximately 6.5 million tons per year (FAO, 2020). The intensification of tilapia farming industry has been confronted with an increasing number of emerging infectious diseases (Ferguson et al., 2014; Eyngor et al., 2014; Machimbirike et al., 2019; Dong et al., 2019; Ramírez-Paredes et al., 2020). For instance, in the past five years, widespread occurrence of disease outbreaks caused by two emerging viruses, namely tilapia lake virus (TiLV) and infectious spleen and kidney necrosis virus (ISKNV), has highlighted the critical need for rapid, accurate diagnostics to aid in the control or mitigation of emerging diseases (Machimbirike et al., 2019; Ramírez-Paredes et al., 2020).

A novel parvovirus, termed *Tilapia parvovirus* (TiPV), was initially discovered from the fecal samples of tilapia-fed crocodiles and intestine samples of tilapia from Hainan, China using next-generation sequencing (Du et al., 2019). However, there was no evidence that this virus was associated with disease in tilapia at that point. Then, Liu et al. (2020) reported their findings of an outbreak investigation involving severe mortality (approx. 60-70%) of adult tilapia (500-600 g) in Hubei, China, since 2015. The diseased fish displayed signs of lethargy, anorexia, abnormal swimming behavior (e.g., corkscrew movements), cutaneous hemorrhages, exophthalmia, and severe ocular lesions on rare occasions (Liu et al., 2020). In this case, although coinfection of TiPV and *Streptococcus agalactiae* was detected in clinically sick fish, TiPV was isolated, propagated in tilapia brain cells (TiB), characterized, and was proved to be pathogenic on its own through experimental infection (reaching ~90% mortalities at 11 dpi).

TiPV is a non-enveloped, spherical virus with a diameter of 30 nm. The TiPV genome contains 4,269 bp linear single-stranded DNA (ssDNA), including 208 bp 5’ UTR, 396 bp ORF1, 1875 bp non-structural protein 1 (NS1), 504 bp NS2, 216 bp ORF2, 1665 bp capsid protein 1 (VP1), and 46 bp 3’ UTR (Liu et al., 2020). TiPV is currently considered a novel *Chapparvovirus* of the *Parvoviridae* family (Du et al., 2019; Liu et al., 2020).

Recently, cases of TiPV infection in hybrid red tilapia were reported in Thailand. The majority of fatalities occurred as a result of complicated coinfection between tilapia lake virus (TiLV) and TiPV, mainly in juveniles (10-30 g), with recorded cumulative mortality rates of 50-75%. Lethargy, hemorrhage, cutaneous ulceration, exophthalmos, and abnormal swimming were all observed in diseased fish (Yamkasem et al., 2021; Piewbang et al., 2022).

To date, pathognomonic lesions serving as the histopathological landmark for diagnosis of TiPV have not been reported. Non-specific histopathological changes reported previously included tissue degeneration/necrosis, inflammatory infiltration by lymphocytes and macrophages, increased numbers of melano-macrophage centers, etc. (Du et al., 2019; Liu et al., 2020; Yamkasem et al., 2021; Piewbang et al., 2022). Here, we report the presence of characteristic Cowdry type A inclusion bodies (CAIB) in the pancreatic tissues during TiPV infection, which may be pathognomonic, and suggest the usefulness of the pancreas as a prime target for histopathological diagnosis of TiPV diseased tilapia. More insights into tissue tropism and localization of TiPV in various infected tissues have also been unveiled.

## Materials and Methods

### Fish samples and preservation

In 2020, we received a set of 10 naturally diseased adult Nile tilapia from an affected breeding farm for disease diagnostics. According to the farm owner, less than 1% of fish in the population showed signs of sickness and only minor mortalities were recorded after handling stress. Freshly dead fish (n=9) were subjected to bacterial isolation, using Tryptic Soy Agar (TSA, Becton, Dickinson, USA) as general medium and *Streptococcus agalactiae* Selective Agar (SSA, HiMedia, India) as a selective medium for *S. agalactiae*. Internal organs (i.e., liver, kidney, spleen, gills, and intestine) from 9 fish were preserved in 10% neutral buffered formalin for histological examination and *in situ* hybridization analysis. The spleen and liver tissues were also preserved in 95% ethanol for molecular diagnostics.

### Histological analysis

Tissue samples were preserved overnight in a 10% neutral buffer formalin solution before being transferred to 70% ethanol for histology. Tissue processing involved dehydration, paraffin embedding, sectioning at 5 μm thickness, and staining with hematoxylin and eosin (H&E). Light microscopy with a digital camera was used to examine histopathological changes.

### Quantitative polymerase chain reaction (qPCR) detection of TiPV and *S. agalactiae*

The preserved spleen and liver samples were subjected to DNA extraction using the TF lysis buffer method (Meemetta et al., 2020). DNA quantity and quality was measured using a NanoDrop (Thermo Scientific, USA), with absorbance set at OD_260_ and OD_280_. DNA samples were diluted to 100 ng/μL for quantitative PCR (qPCR) reactions. Detection of TiPV was performed using a protocol described by Liu et al. (2020). PCR mixtures were comprised of 10 μL 2X SYBR Green Master Mix (Bio-Rad, USA), 0.5 μL of each forward and reverse primers (10 μM of each TiPV-Fq/TiPV-Rq) (Table 1), 2 μL DNA template and 7 μL ddH_2_O. PCR was performed at 94 °C for 2 min followed by 40 cycles of 94 °C for 10 s and 60 °C for 30 s. The melting curve was analyzed in the last cycle. On the other hand, probe-based qPCR targeting *groEL* gene was used to diagnose *S. agalactiae* infection, according to Leigh et al. (2019). The qPCR reagents mixture contained 10 μL of 2X iTaq Universal Probes Supermix (Bio-Rad, USA), 0.5 μL of 10 μM SagroEL-probe (Table 1), 1.8 μL of each primer SagroEL-F/R (10 μM), 3 μl of DNA template and 2.9 μL ddH_2_O. The qPCR reaction was performed at 95 °C for 2 min followed by 40 cycles of 95 °C for 5 s and 60 °C for 30 s. DNA extracted from *S. agalactiae* 2809 (Linh et al., 2021) was used as a positive control, and ddH_2_O without DNA template was used as a negative control.

**Table 1:**
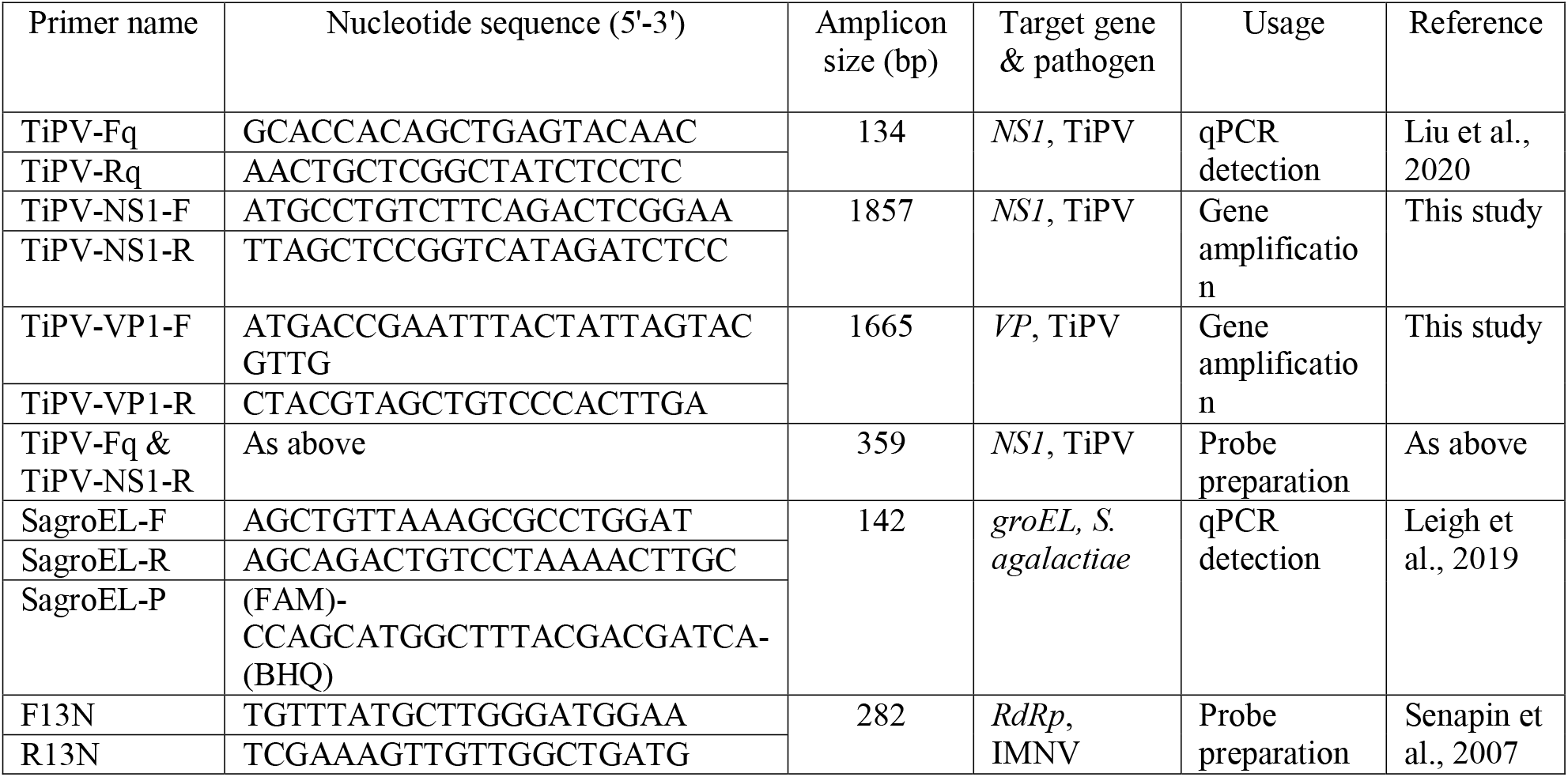
Primers used in this study

### Amplification and sequence analysis of *NS1* and *VP1* genes of TiPV

For confirmation of TiPV, full-length *NS1* (1857 bp) and *VP1* (1665 bp) were amplified using newly designed primers, TiPV-NS1F/R and TiPV-VP1F/R (Table 1), respectively. The primers were designed based on the complete genome sequence of TiPV (accession number MT393593). Reaction mixtures comprised of 2 μL 10X PCR buffer, 0.4 μL of 10 mM dNTP, 0.4 μL of 50 mM MgCl_2_, 0.5 μL of each forward and reverse primers (10 μM), 0.1 μL Taq polymerase (Invitrogen, 5U/μL), 2 μL DNA template, and 14.1 μL ddH_2_O. The PCR protocol started with a pre-denaturation at 95 °C for 5 min followed by 30 cycles of denaturation at 95 °C for 30 s, annealing at 60°C (for *NS1*) or at 54 °C (for *VP1*) for 30 s, extension at 72 °C for 2 min, and a final extension at 72 °C for 5 min. PCR products were visualized on 1% agarose gel, and expected amplicons of *NS1* and *VP1* genes were purified using Nucleospin^®^ Gel and PCR clean-up kit (Takara Bio, Japan). Purified PCR amplicons were individually subjected to Barcode-Tagged sequencing (Celemics, Inc., South Korea). The obtained DNA sequences were processed by Geneious Prime software (Biomatters, New Zealand) and analyzed using BLASTn from NCBI database.

### *In situ* hybridization (ISH)

Localization of TiPV in the infected tissues was investigated using an ISH assay. TiPV-specific digoxygenin (DIG) labeling probe (359 bp) was synthesized using primers TiPV-Fq and TiPV-NS1-R (Table 1) and DIG-labeling mix reagent (Roche, Germany) according to the manufacturer’s instructions. The PCR protocol included 30 cycles: pre-denature 94 °C for 5 min, denature 94 °C for 30 s, annealing 55 °C for 30 s, extension 72 °C for 30 s, final extension 72 °C for 2 min. The DNA-labeling probe was gel purified using Nucleospin^®^ Gel and PCR clean-up kit. An unrelated probe derived from a 282 bp fragment of shrimp infectious myonecrosis virus (IMNV) (Senapin et al., 2007) was used as a negative control. The standard ISH procedure was conducted with a probe concentration of 500 ng/slide (Dinh-Hung et al., 2021). Three consecutive sections from each sample were assayed using a TiPV-specific probe, unrelated probe, and H&E stain. The results were interpreted in parallel under a light microscope connected to a digital camera.

## Results

### Gross signs and histopathology

Grossly, the fish submitted to our laboratory presented with loss of body condition, skin erosions, cloudy eye lenses, and corneal injury (Figure 1A, B). Internal organs exhibited no obvious abnormal changes in most individuals, except for rare instances of microhepatica, intestinal pallor, and necrotic areas in the ovary (Figure 1C).

**Figure 1.**
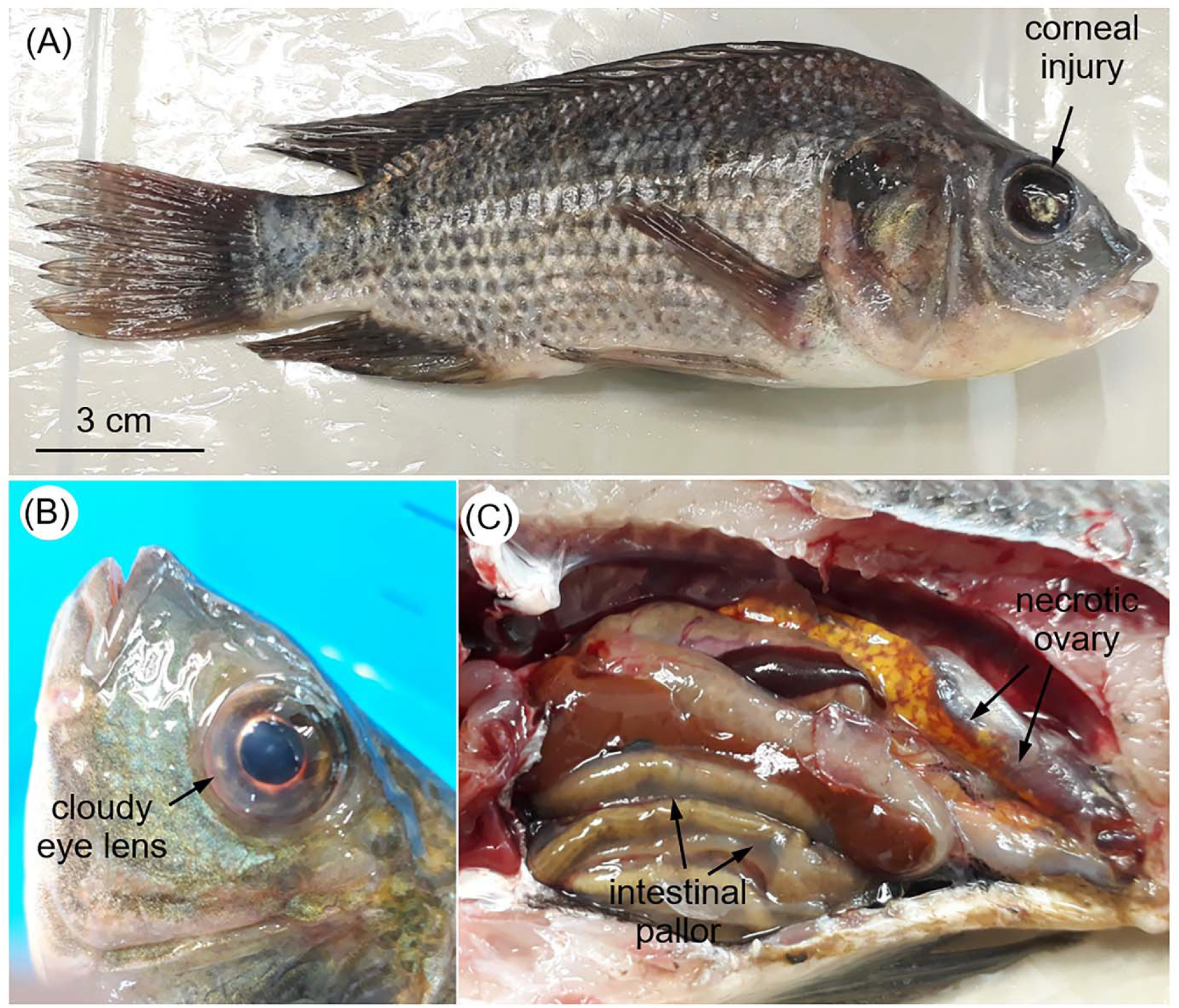
Gross signs of Nile tilapia naturally infected with TiPV showing corneal injury (A), cloudy eye lens (B), liver atrophy, intestinal pallor and necrotic ovary (C).

Histopathologically, Cowdry type A inclusion bodies (CAIB) in pancreatic acinar cells (in both pancreatic islands of the hepatopancreas and pancreatic tissue along the intestine) were recorded in all fish (Table 2). CAIB are intranuclear inclusions distinguished by chromatin margination and a distinct, clear halo surrounding a central basophilic inclusion body (Figure 2A-D). There were differences between CAIB size, ranging from small inclusions (early stage) similar to nucleoli to eccentric, large inclusions with a well-defined halo (late stage) (Figure 2E).

**Table 2:**
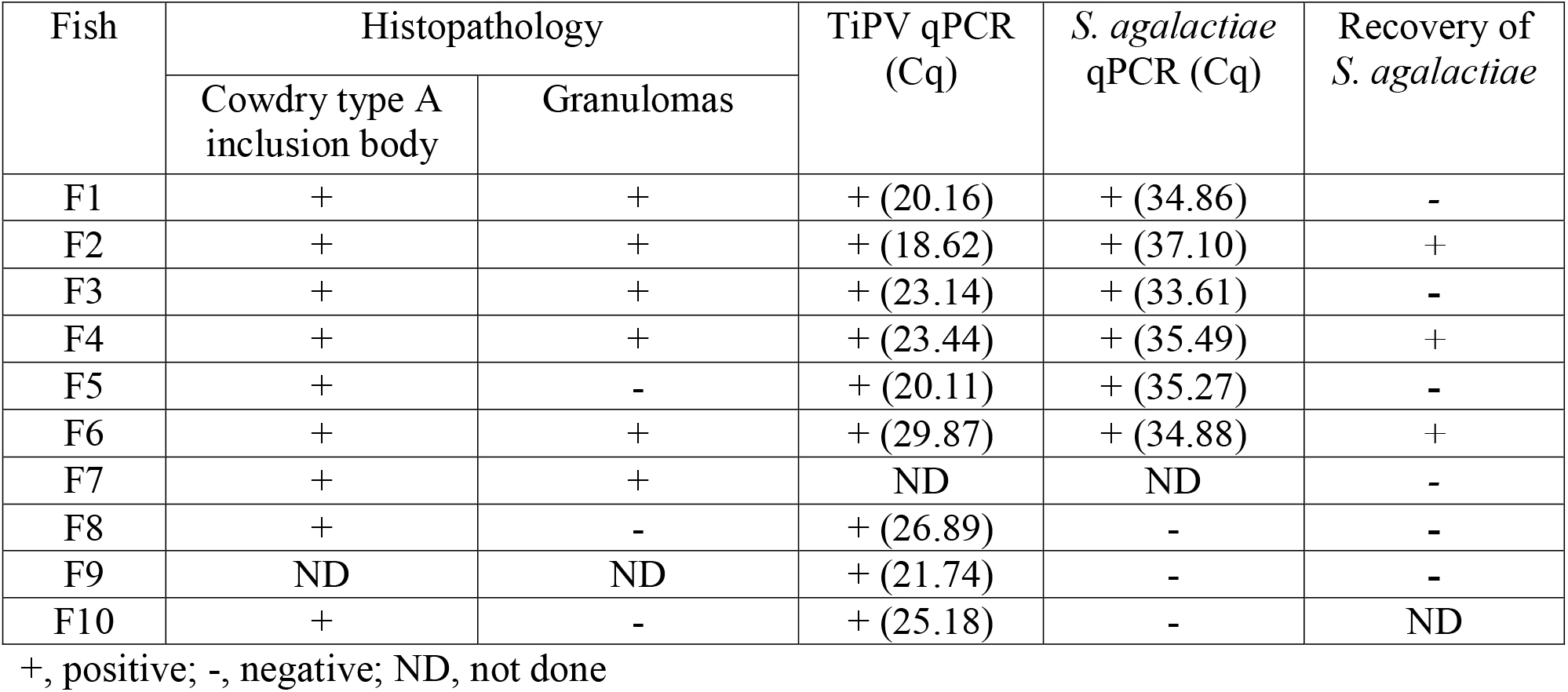
Summary of diagnostic results

| Fish | Histopathology |  | TiPV qPCR (Cq) | *S. agalactiae* qPCR (Cq) | Recovery of *S. agalactiae* |
| --- | --- | --- | --- | --- | --- |
|  | Cowdry type A inclusion body | Granulomas |  |  |  |
| F1 | + | + | + (20.16) | + (34.86) | - |
| F2 | + | + | + (18.62) | + (37.10) | + |
| F3 | + | + | + (23.14) | + (33.61) | - |
| F4 | + | + | + (23.44) | + (35.49) | + |
| F5 | + | - | + (20.11) | + (35.27) | - |
| F6 | + | + | + (29.87) | + (34.88) | + |
| F7 | + | + | ND | ND | - |
| F8 | + | - | + (26.89) | - | - |
| F9 | ND | ND | + (21.74) | - | - |
| F10 | + | - | + (25.18) | - | ND |

+, positive; -, negative; ND, not done

**Figure 2.**
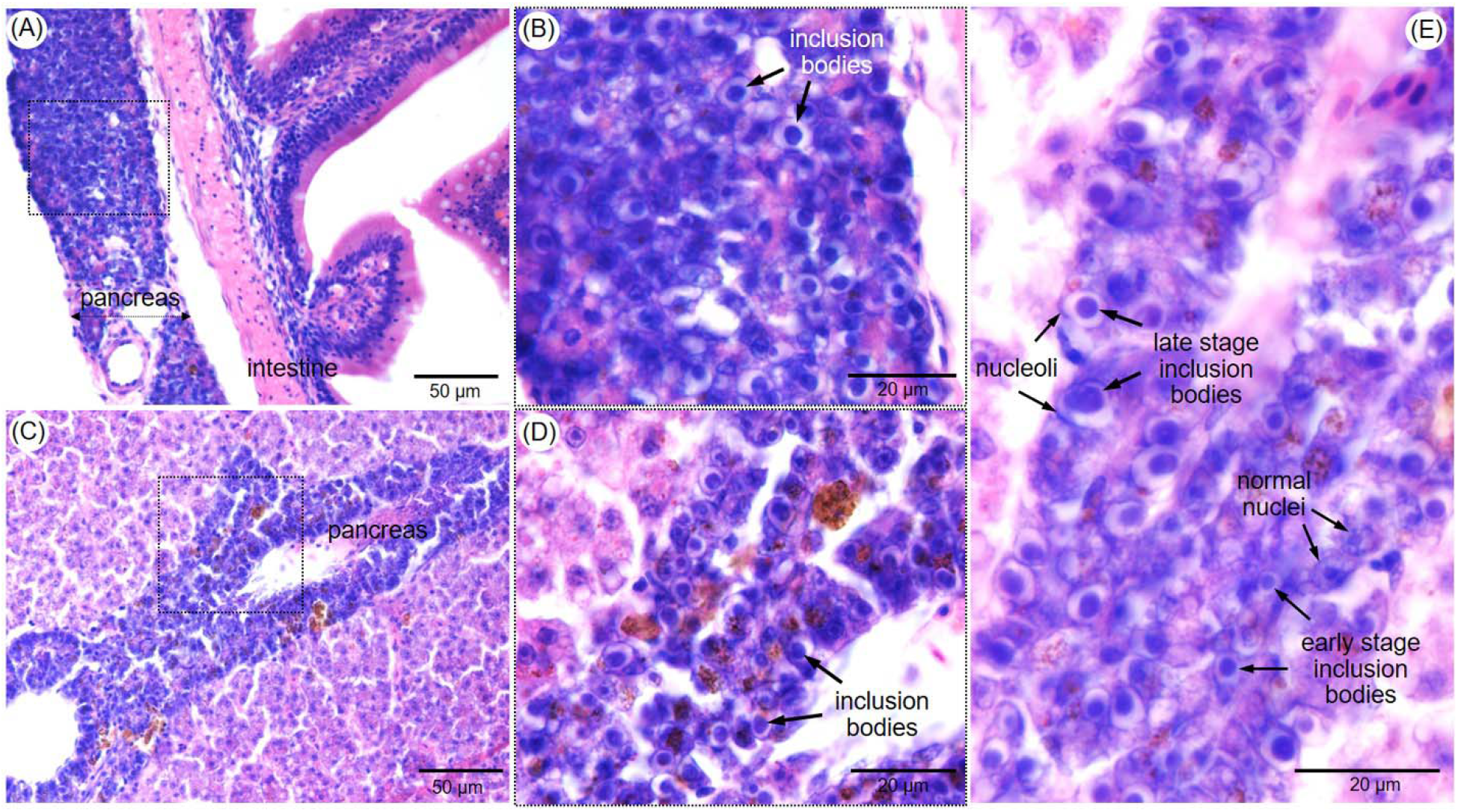
Microphotographs of H&E-stained sections of the pancreatic tissue along the intestine (A-B) and within the hepatopancreas (C-E) of clinically sick tilapia infected with TiPV. Pictures B and D are higher magnification images of the areas marked with dotted boxes in A and C, respectively. Typical Cowdry type A inclusion bodies are visualized in B and D. Early and late-stage inclusion bodies are clearly observed in the pancreatic cells (E).

Additionally, characteristic granulomas associated with intralesional bacterial cocci were observed in the majority of the fish examined, most notably in the spleen, kidney, brain, and ovary (Figure 3A-D). Typically, granulomas were filled with dark-brown pigment (consistent with melanin). Numerous cocci bacteria (morphologically consistent with *Streptococcus* sp.) were observed at high magnification in the necrotic core of some granulomas (Figure 3E) or intracellularly on the membrane of oocytes (Figure 3F).

**Figure 3.**
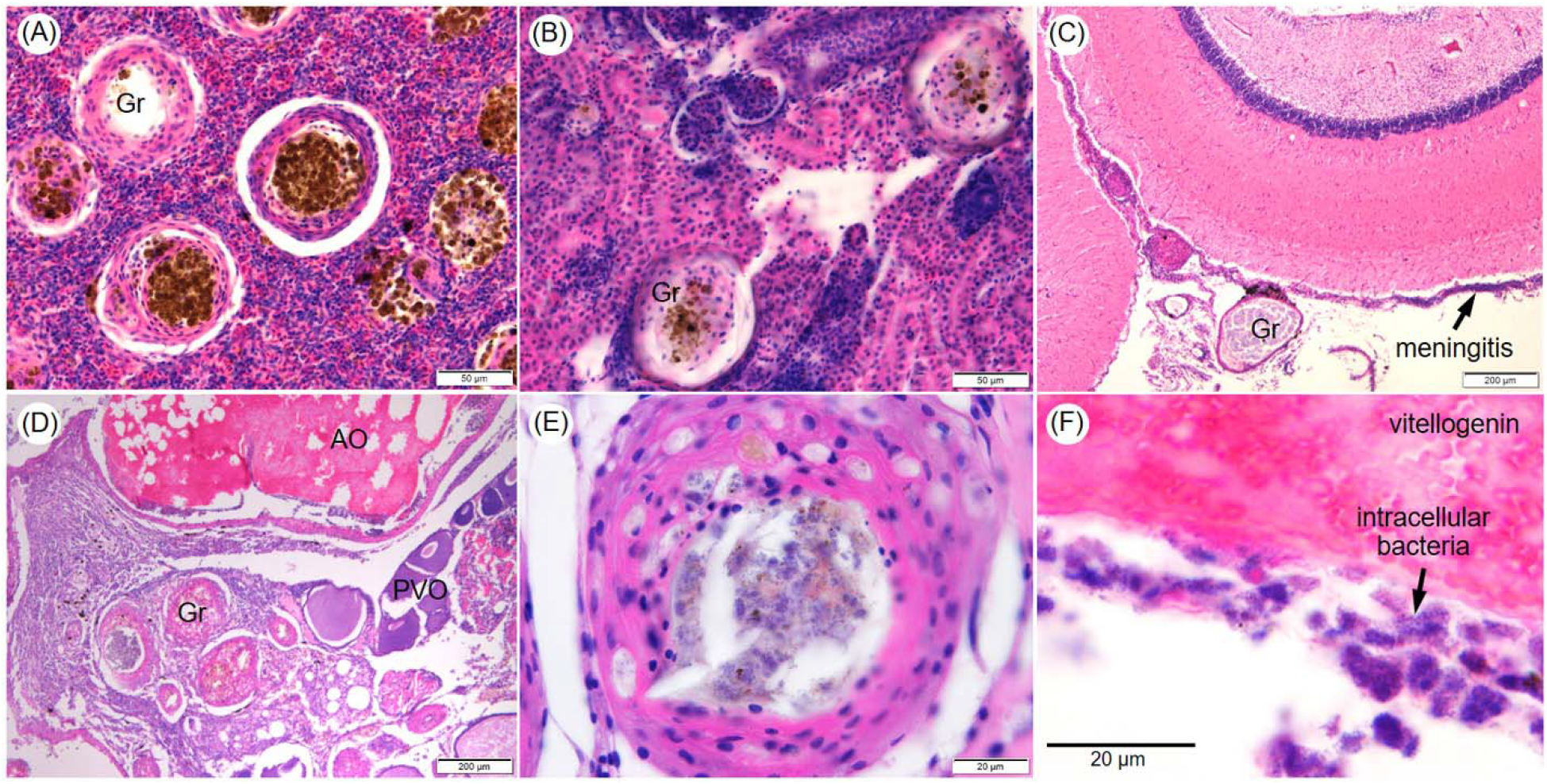
Representative microphotographs of the spleen (A), kidney (B), brain (C) and ovary (D) sections of a clinically sick tilapia showing presence of granulomas containing intralesional pigment (melanin), and increased melanomacrophage centers. Severe meningitis was observed in the brain (arrow, C). High magnification of a typical granuloma showing numerous bacterial cocci in the center (E). Numerous intracellular bacterial cocci on the membrane of oocytes in the ovary (arrow, F). AO, atretic oocyte; PVO, pre-vitellogenic oocytes; Gr, granuloma. H&E staining.

### Diseased fish tested positive for both TiPV and *S. agalactiae*

All examined fish (n = 9) were tested positive for TiPV by qPCR with Cq values ranging from 18.62 to 29.87 (Table 2). Subsequently, *NS1* and *VP1* of TiPV were successfully amplified from representative samples and sequenced. Their partial ORF sequences of 1803 and 1414 bp were submitted to GenBank under accession numbers OM884999 and OM885000, respectively. Blast searches revealed that the *NS1* and *VP1* in this study have 99.5 and 99.22% nucleotide identities to the *NS1* and *VP1* of the reference China TiPV strain TiPVC20160912 (accession no. MT393593.1), respectively. Compared to the Thai TiPV strain KU01-TH/2020, the percent nucleotide identity was 99.72% for the *NS1* (accession no. MW685502.1) and 99.58% for the *VP1* (accession no. MW685502.1), respectively.

Six out of nine samples tested positive for *S. agalactiae* using *S. agalactiae* specific qPCR with Cq ranging from 33.61 to 37.10. Pinpoint colony-forming bacteria were successfully cultured from 3/9 fish, and all tested positive for *S. agalactiae* by qPCR (Table 1).

### Localization of TiPV in the infected tissue by ISH

Using a DIG-labeled probe targeting a 359-bp fragment of the *NS1* gene of TiPV, there were positive signals in multiple organs of TiPV infected fish, including the liver and pancreas (Figure 4) as well as the kidney, spleen, gills, and ovary (Figure 5). In contrast, no positive signal was found when the same samples were assayed with an unrelated probe (negative control - Figure 4–5). At the cellular level, TiPV positive signals were clearly localized to the nuclei and were noted in hepatocytes (Figure 4A-C), pancreatic acinar cells (Figure 4D-F), renal tubular epithelial cells (Figure 5A-C), splenic ellipsoids (Figure 5D-F), and cells within primary and secondary gill lamella (Figure 5G-I). Notably, positive signals were also detected in cells around the oocyte cell membrane in the ovary (Figure 5 J-L). CAIB were devoid of ISH signal (data not shown).

**Figure 4.**
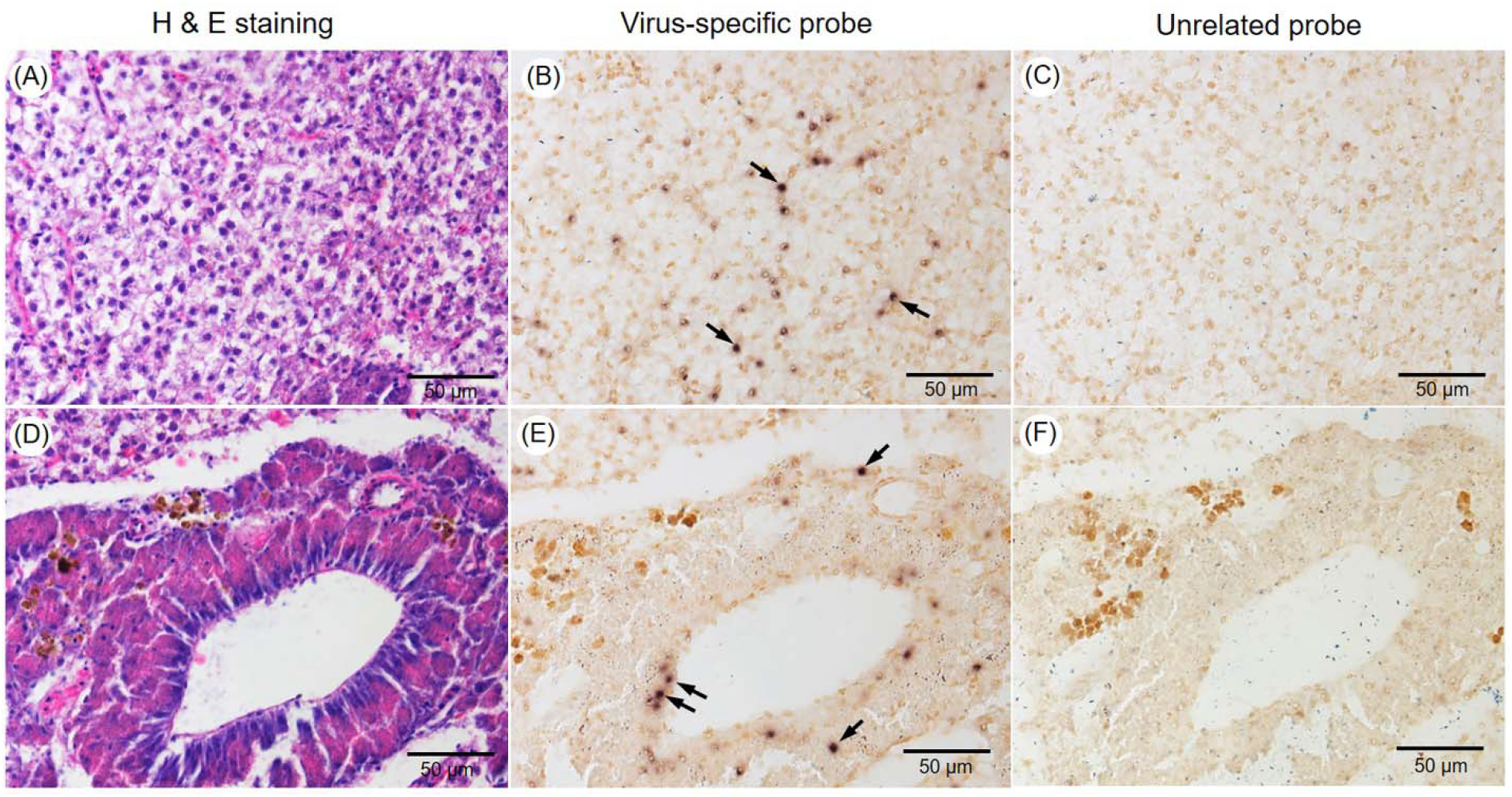
Representative photomicrographs of three continuous sections of the hepatopancreas (A-C) and pancreas (D-F) of a TiPV infected tilapia. H&E-stained sections (A, D), ISH with TiPV-specific probe (B, E), and ISH with unrelated probe (C, F – negative control). A positive signal is characterized by intranuclear dark brown staining, which is multifocal in both hepatocytes and pancreas (arrows in B and E).

**Figure 5.**
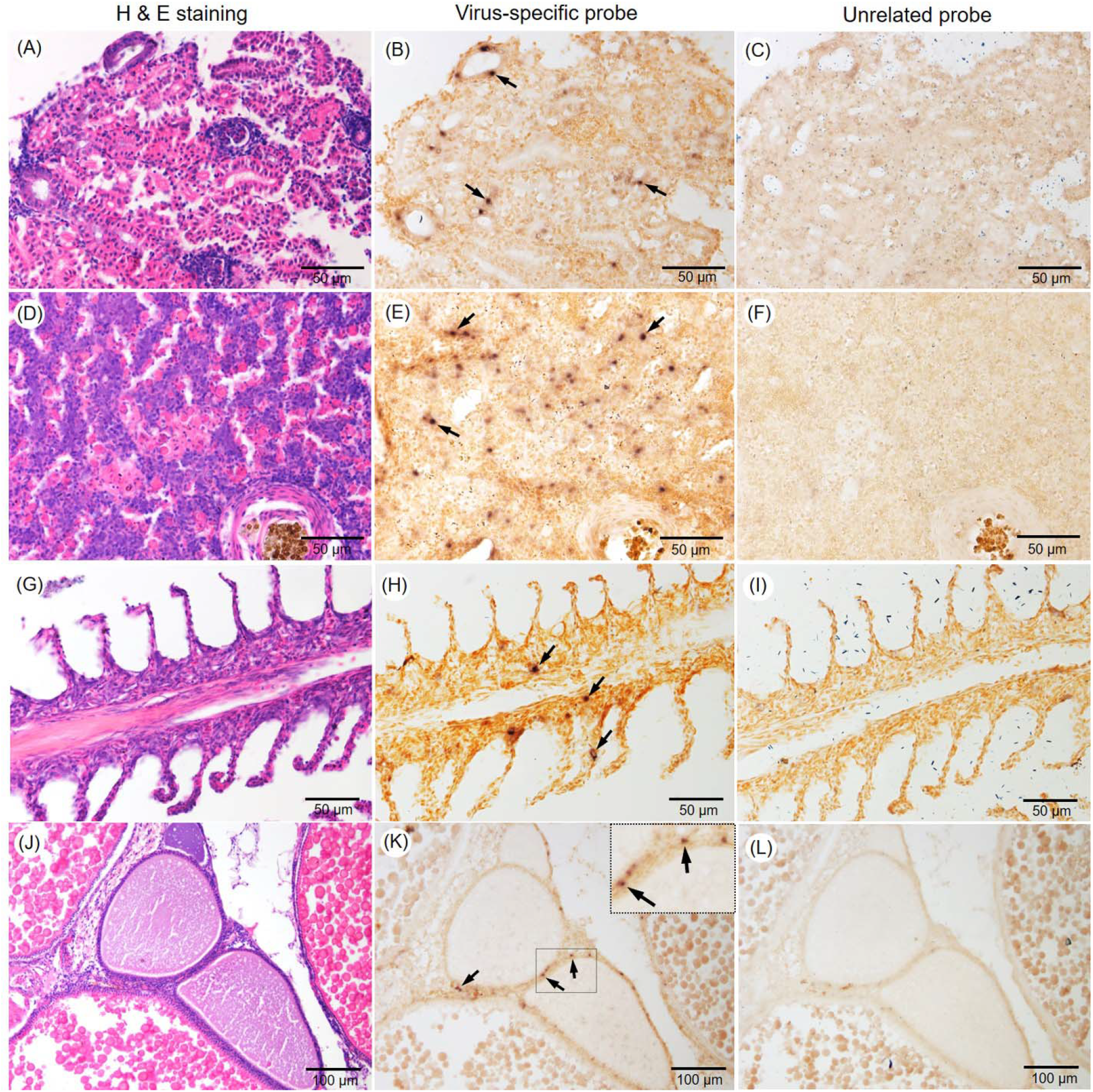
Representative photomicrographs of three continuous sections of kidney (A-C), spleen (D-F), gills (G-I), and ovary (J-L) of TiPV infected tilapia. H&E-stained sections (A, D, G and J), ISH with TiPV-specific probe (B, E, H and K), and ISH with unrelated probe (C, F, I and L). TiPV positive signals are indicated by intranuclear dark brown staining (arrows).

## Discussion

This study reported the presence of CAIB in the pancreas of TiPV-infected tilapia for the first time. This pathological feature was previously identified to be a histopathological landmark for the diagnosis of two parvovirus infections in shrimp, namely hepatopancreatic parvovirus (HPV) (recently called decapod hepanhamaparvovirus, DHPV) and infectious hypodermal and haematopoietic necrosis virus (IHHNV) (Flegel et al., 2006; Srisala et al., 2021) and terrestrial species (e.g. cattle, dogs, cats) (Kennedy & Palmer, 2016). Interestingly, the CAIB observed in this study are similar to HPV inclusion bodies in shrimp, but the CAIB associated with TiPV infection appear to be less numerous and smaller, requiring observation under high magnification (100X). Although general histopathological changes have been reported, this previously overlooked inclusion body is particularly useful for diagnosing TiPV due to its characteristic morphology. Furthermore, CAIB were detected in the pancreas but not in other infected tissues, demonstrating the pancreas’ utility as a primary target for histopathological diagnosis of TiPV infection in tilapia. This finding is important for identifying TiPV in new geographic locations, which typically requires a combination of macroscopic, microscopic, and molecular investigation. Also, this unique histopathological characteristic enables retrospective investigation of TiPV infection in archived samples. It is possible to hypothesize that the pancreas may present unique features in the pathogenesis and host-pathogen interaction of TiPV infection in tilapia.

Although ISH with a TiPV-specific probe detected viral genomes in the nuclei of the cells from multiple organs in TiPV infected tilapia, no hybridization signal was detected within CAIB. It is possible that this is due to a defective replication cycle leading to aberrant accumulation of virions lacking nucleic acid, or that the thickness of viral protein that makes up the inclusion bodies prevents TiPV probe penetration. Interestingly, this phenomenon has also been observed in shrimp parvovirus (per. comm., Prof. Tim W. Flegel), and warrants further investigation. Similar to the previous reports (Liu et al., 2020; Piewbang et al., 2022), TiPV was detected in various organs and cell types, indicating widespread cellular tropism, and the capability of infecting tilapia systemically. This finding is interesting, as parvoviruses are reported to be radiomimetic, i.e. they have tropism for dividing cells in other species (Boes and Durham, 2017), a feature that would be expected to also occur in TiPV, and may suggest that division is ongoing in adult tilapia in all of the tissues targeted by TiPV. Notably, the presence of TiPV positive signals in oocyte membranes may indicate that this virus is capable of vertical transmission similar to that of TiLV (Dong et al., 2020). However, more research would be necessary to evaluate TiPV vertical transmission.

Although ISH was not done for *S. agalactiae* detection, the presence of dense intracellular cocci bacteria was also occasionally observed in the epithelial membrane of oocytes in the degenerated ovary of the affected female fish, providing supporting evidence for vertical transmission of this agent (Pradeep et al., 2016).

The high nucleotide sequence similarity between the TiPV isolates in this study and those reported previously in China and Thailand (Du et al., 2019; Liu et al., 2020; Yamkasem et al., 2021) suggests that they may be from the same clone or have a highly conserved genomic nature. Further genomic investigations could provide insight into viral epidemiology and evolution.

The disease event in this study could be attributed to TiPV and *S. agalactiae* coinfection. Similar to a previous report, TiPV and *S. agalactiae* coinfection was associated with a disease outbreak in adult tilapia in China. TiPV was also found in a relatively high prevalence (23.1 to 64.6 %) of randomly collected samples without clinical signs in multiple geographical locations in China (Liu et al. 2020). Most recently, TiPV and TiLV coinfections were found in the majority of disease cases in juvenile tilapia in Thailand (Yamkasem et al., 2021; Piewbang et al., 2022). Despite the fact that the fish population in this study was infected with both TiPV and *S. agalactiae*, only a small proportion of mortality was observed after handling stress, implying that both pathogens were in the chronic phase of infection. With respect to *S. agalactiae*, the presence of numerous granulomas is marked as a histopathological feature of chronic infection. Taken together, the available evidence indicates that additional research into the pathogenicity of TiPV and risk factors for disease outbreaks is necessary to determine whether TiPV is a true pathogen or an opportunistic agent. As TiPV has been described recently, its pathogenesis is unclear. However, in view of this coinfection, it is possible to hypothesize that its presence may lead to immunosuppression and facilitate secondary infection. For example, such an effect has been reported in canine parvovirus type 2, feline panleukopaenia virus, and bovine viral diarrhea virus. In these instances, immunosuppression is interpreted as a direct result of lymphocytolysis leading to lymphoid atrophy (Boes and Durham, 2017).

In conclusion, the present study described the presence of CAIB in the pancreas as a histopathological landmark for the diagnosis of TiPV in tilapia. This pathological lesion is unique to the pancreas, highlighting the critical nature of this tissue as a prime target for histopathological diagnosis. Initial detection of CAIB is particularly helpful to suggest the need for PCR, sequencing, and ISH analysis to identify TiPV infection. This is especially relevant for TiPV diagnosis in countries or regions where it is reported for the first time, in order to facilitate rapid diagnosis and emergent response to mitigate TiPV’s negative impact on tilapia aquaculture.

## Acknowledgements

This study was supported by BIOTEC Fellow’s Research Grants (P-19-50170). H.T. Dong acknowledges the Research Initiation Grant from the Asian Institute of Technology.

## Declaration of Competing Interest

The authors declare that they have no known competing financial interests or personal relationships that could have appeared to influence the work reported in this paper.

## Credit authorship contribution statement

H.T.D., P.S., and S.S., conceptualization, methodology, investigation, writing original draft; L.T.D., T.T.M., N.V.L., investigation, J.D.P., K.R.S., review-editing. All authors have read and agreed to the current version of the manuscript.

## Notes

### Competing Interest Statement

The authors have declared no competing interest.

